# Diabetes mellitus affects urinary tissue tropism of group B streptococci in a sex-dependent manner

**DOI:** 10.1101/2025.05.02.651940

**Authors:** Preeti P John, Bidhan Gautam, Kenneth A Rogers, Ritwij Kulkarni

## Abstract

Diabetes mellitus (DM) increases susceptibility to *Streptococcus agalactiae* (group B *Streptococcus* or GBS) urinary tract infections (UTI) and their exacerbations such as ascending pyelonephritis and sepsis, although underlying molecular mechanisms have not been fully deciphered. To address our hypothesis that DM mediated alterations in the host urinary immune defenses increase susceptibility to GBS-UTI, we inoculated, via transurethral route, GBS strain 10/84 into the urinary bladders of 8 weeks old, male and female, diabetic (obese, hyperglycemic mice denoted as D) and non-diabetic (ND) mouse littermates. At 24 h post-infection (hpi), compared to their ND littermates, the D-mice showed significantly higher GBS CFUs in the bladder, the kidneys, and the spleen. This was accompanied by a significantly reduced recruitment of CD45^+^ leukocytes to and suppression of pro-inflammatory cytokine production in GBS-infected D-bladder and D-kidneys compared to their ND counterparts.

We also noted sex-dependent differences in GBS urinary tissue tropism. For example, D-females exhibited significantly higher bladder burden compared to ND-females, D-males, and to ND-males, while the bladder burden in ND-females was not significantly different from that in ND-males. In contrast, ND-males showed significantly higher kidney CFUs than ND-females and DM further increased kidney CFU burden in males. GBS-induced recruitment of CD45^+^ leukocytes was not different between male and female mice within either D or ND cohorts. GBS-UTI induced significantly higher CXCL1 production in male bladders in both D and ND cohorts compared to their female counterparts.

These results indicate that DM increases susceptibility to GBS-UTI and the dissemination to spleen by affecting leukocyte recruitment and pro-inflammatory cytokine production in a sex-dependent manner.

**Importance:** In this study we sought to understand why diabetic individuals are more susceptible to urinary tract infections (UTI) by Gram positive *Streptococcus agalactiae* (group B *Streptococcus* or GBS). We induced UTI by inoculating GBS into the bladders of diabetic (obese, hyperglycemic) and non-diabetic, male and female mice. At 24 hours after infection, compared to the non-diabetic mice, the diabetic mouse urinary tracts showed higher GBS counts and reduced immune defenses. Diabetes also promoted dissemination of GBS to the spleen. Furthermore, female diabetic mice were more susceptible to bladder infection while diabetes worsened the increased overall susceptibility of males to kidney infection, a difficult-to-treat, potentially life threatening exacerbation of GBS-UTI. Overall, our results suggest that both diabetes and sex are important determinants of susceptibility to GBS-UTI and resulting severe outcomes.

## INTRODUCTION

*Streptococcus agalactiae* (group B *Streptococcus* or GBS) is an asymptomatic colonizer of human gastrointestinal and genitourinary tracts and an atypical uropathogen that causes an estimated 3-4.5 million urinary tract (UT) infections (UTI) globally including 160,000 cases per year in the US (1-4). Diabetes mellitus (DM) increases the risk of GBS asymptomatic bacteriuria, cystitis, and exacerbations including pyelonephritis and bloodstream infections (BSI) according to some (4-11) but not all (12) published reports. The primary objective of this study was to identify specific anti-GBS urinary immune defenses affected by DM and bridge the knowledge gap that the mechanisms underlying the increased susceptibility of diabetic individuals to GBS UTI have not been fully deciphered.

Various host factors unique to the diabetic urinary microenvironment have been implicated in increasing the risk of UTI in diabetic individuals. For example, glycosuria and urine retention due to neurogenic bladder may enhance the growth of uropathogenic bacteria against which immune defenses weakened by DM, such as reduced phagocytosis, intracellular killing, may be ineffective (13-15). We have previously reported that *in vitro exposure* to glycosuria significantly augments both growth as well as virulence of GBS (16). Patras *et al* have reported that significantly higher GBS burden in the urinary bladders of streptozotocin (STZ)-induced diabetic mice is associated with higher levels of antimicrobial peptide cathelicidin (LL-37) and increased mast cell activity, although both these immune defenses appear to be ineffective against GBS in diabetic as well as non-diabetic murine UT (17). STZ, a DNA alkylating glucose analog, that destroys endogenous beta cells, resulting in reduced endogenous insulin production and hyperglycemia, is a versatile chemical that can be used to induce DM in a variety of animal models (18). Indeed, researchers have effectively used the mouse model of STZ-induced DM to study the immunopathology of ascending UTI by uropathogenic *Escherichia coli* (UPEC), *Klebsiella pneumoniae*, or *Enterococcus faecalis* (19-21). However, STZ treatment causes serious, adverse side effects including nephrotoxicity and immunosuppression (lymphopenia) which can confound the analysis of urinary immune responses in diabetes (22, 23). Moreover, the marked resistance of female mice to the diabetogenic effects of STZ necessitates administration of higher, potentially toxic level, doses of STZ for inducing DM in female mice (24), in turn making the mouse model of STZ-induced DM unsuitable for studying sex-based differences in the pathophysiology of diabetic UTI. Hence, in this study we used the *db/db* mouse model of genetically induced DM which models human type 2 DM associated with obesity and hyperglycemia (25). The *db/db* mouse model of DM has been previously used to reveal the role of insulin signaling in renal antibacterial defenses against ascending UTI caused by UPEC (26).

The transurethral catheterization of GBS 10/84 into the urinary bladders of 8 weeks old, *db*/*db* diabetic (D) and non-diabetic (ND) littermates revealed that DM increases the overall susceptibility to UTI as well as uBSI. Moreover, DM also affected GBS organ tropism in a sex-dependent manner. Thus, at 24 h post-infection (hpi), compared to their ND counterparts, D-females were more suscpetible to GBS-cystitis, D-males were more suscpetible to GBS-pyelonpehritis, while both D-male and D-female mice showed signficiantly higher spleen GBS burden. We also noted signficiantly reduced recruitment of CD45^+^ total leukocytes to GBS-infected D-bladder and D-kidneys and significant reduction in CXCL1 in GBS-infected D-bladders and in CCL3, IL-1β, and IL-17 in GBS-infected D-kidneys compared to their ND counterparts. Based on these results we conclude that (i) DM increases susceptibility to not only GBS-UTI but also GBS dissemination to the spleen (ii) which is driven by reduced leukocyte recruitment to and lower cytokine production in GBS-infected UT and (iii) confluence of male sex and diabetes induces the ascent of GBS to kidneys resulting in pyelonephritis.

## Materials and Methods

### Model uropathogen

*Streptococcus agalactiae* CNCTC 10/84 mid-log phase (OD600 = 0.6) culture stocks stored in 20% glycerol-sterile tryptic soy broth (TSB) at −20°C were thawed, washed in sterile phosphate-buffered saline (PBS), resuspended in sterile tryptic soy (TS) broth, and incubated at 37°C to mid-log phase before use in experiment.

### The murine model of ascending UTI in diabetes

Protocols for mouse husbandry and experimentation were approved by IACUC at UL Lafayette (#2020-8717-025-VLW). The breeding trios of heterozygous mice from BKS.Cg-*Dock7*^*m*^ +/+ *Lepr*^*db*^/J (Jackson Lab strain #000642 referred to as Dock7) were maintained in the vivarium at UL Lafayette to generate the male and female, diabetic (*Lepr*^*db/db*^ denoted as D) and non-diabetic (ND) littermates. Mice were genotyped at 5 weeks of age. Mice were weighed and their blood glucose levels were monitored once a week from 5 weeks of age until infection. In addition, D-mice from B6.BKS(D)-*Lepr*^*db*^/J background (Jackson Lab strain #000697 referred to as BKS) were examined for GBS organ burden. Mice were infected via transurethral catheterization with 10^7^ CFU of GBS10/84 in 50μl PBS. At 24hpi, mice were euthanized. To avoid cross-contamination from dissection tools, the spleen, the most distal organ from the site of GBS inoculation, was harvested first, followed by the kidneys. The urinary bladder, the site of GBS inoculation, was harvested the last. The organ homogenates in sterile PBS were processed for CFU burden by dilution plating, immune cell recruitment by flow cytometry, or cytokine levels by ELISA.

The spread for organ burden data was similar for the D-mice from Dock7 and BKS backgrounds, hence the CFU data from these mouse strains were analyzed and presented together in Fig 1. Only the Dock7 littermates were used for flow cytometry (Fig 2) and ELISA (Fig 3).

**FIG 1.**
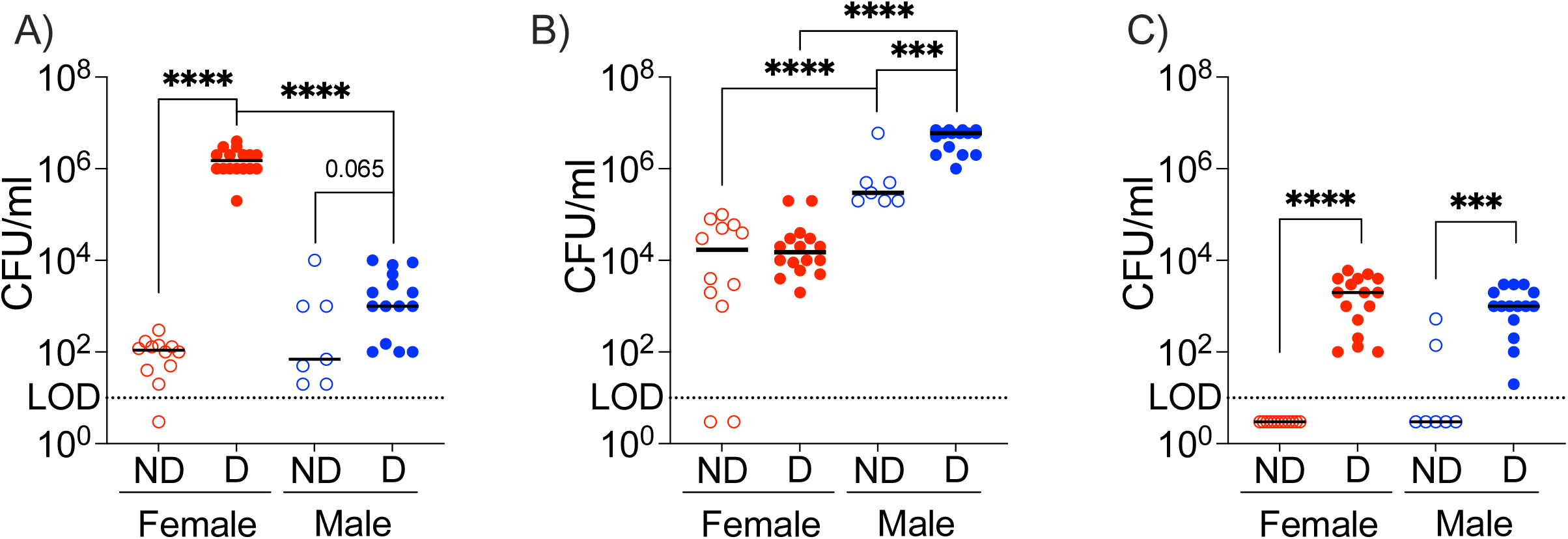

**FIG 2.**
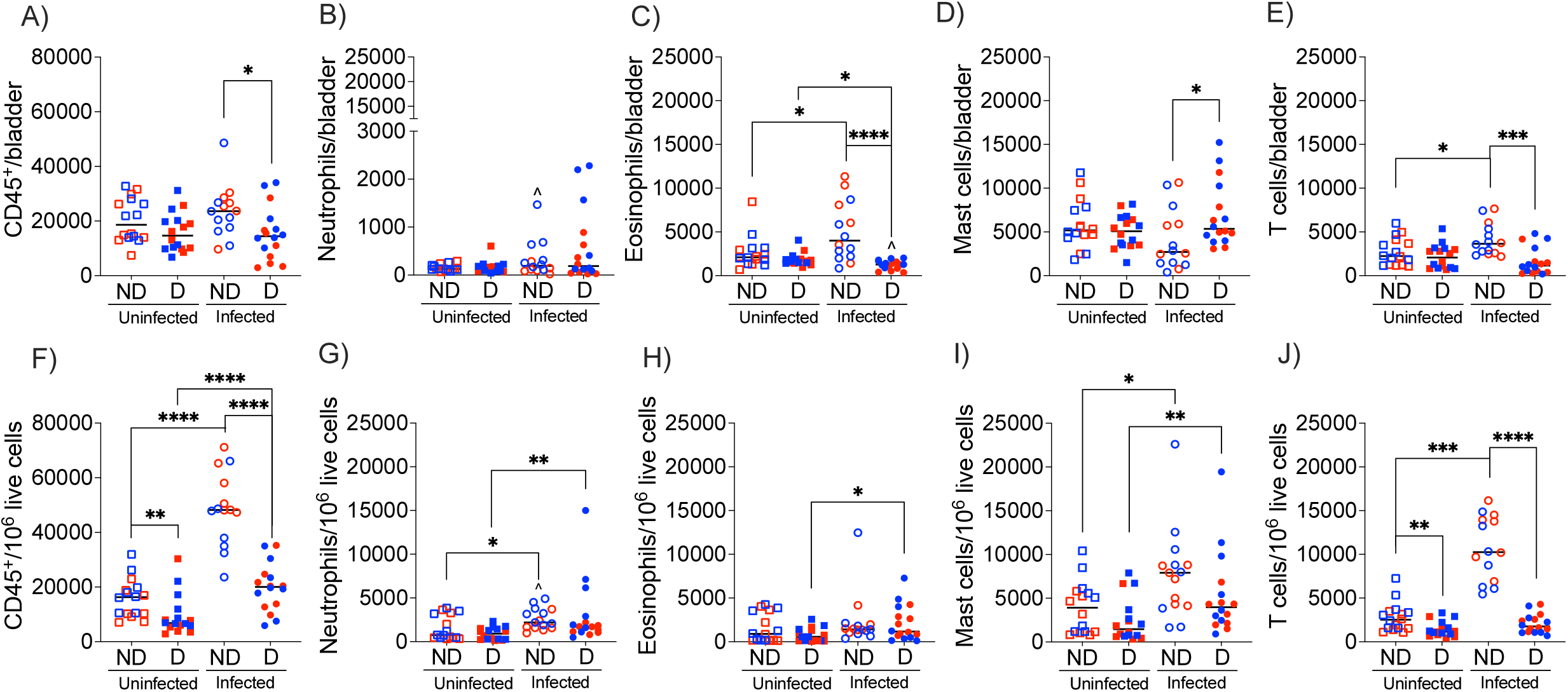

**FIG 3.**
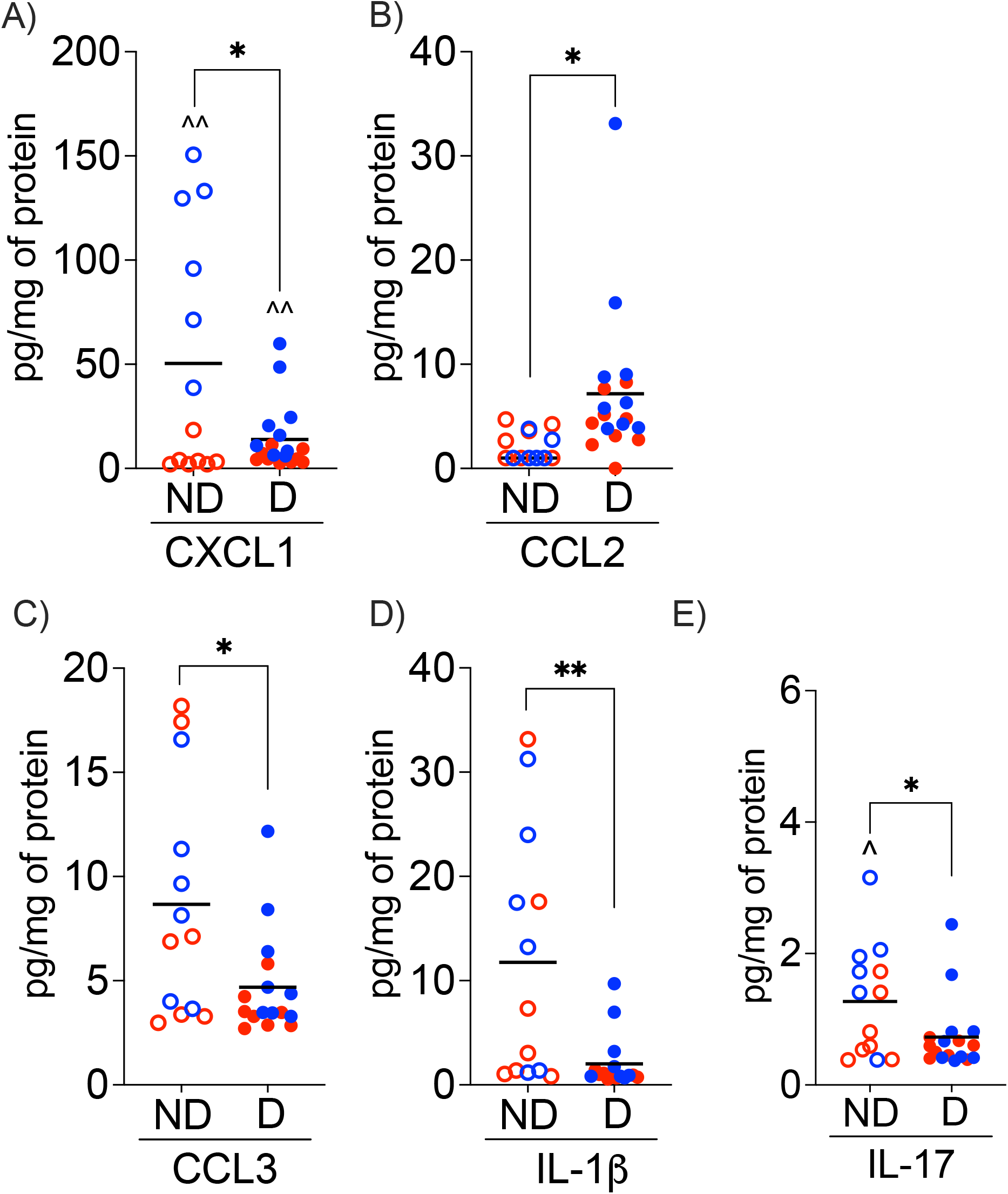

### Bacterial organ burden

Homogenized bladder, kidney, and spleen tissues from different mouse cohorts collected at 24 hpi were dilution plated on blood agar plates and incubated overnight at 37°C. The bacterial organ burden from each mouse is presented as a scatter plot with median as the measure of central tendency.

### Flow cytometry analysis

Whole bladders and sections of kidney tissue were minced followed by enzymatic digestion in RPMI medium containing collagenase IV (10 mg/ml for the bladder and 2 mg/ml for the kidney) and DNase I at room temperature for 90 min and 250 rpm shaking with frequent pipetting to mix. The cell suspension was passed through a 35 μm strainer (Falcon®) to filter cell debris and washed once in D-PBS (2000 rpm, 5 min, RT). After treatment with RBC lysis buffer (RT, 10 min), the cells were centrifuged. Next, the cell pellets were stained (in tubes protected from light) with 1 μl live/dead marker (Alexa Fluor 430 NHS Ester (Succinimidyl Ester), ThermoFisher) at RT for 25 min. This was followed by surface staining first with 2 μl of Fc block (4°C for 10 min) and then with a cocktail of fluorescent-labelled antibodies (2 μl/antibody) to identify specific immune cells (RT for 15 min) (17). The cell pellets were washed once in FACS buffer (D-PBS + 2%FBS) between staining steps. After the last step, cells were incubated in 250 μl fixation buffer at 4°C for 20 min, washed once in FACS buffer, and resuspended in FACS buffer for use in flow-cytometry. The data were analyzed with FlowJO ^™^ version 10. After gating on CD45^+^ total leukocytes, MHCII^-^CD11b^+^Ly6G^+^ were regarded as neutrophils, MHCII^-^CD11b^+^SiglecF^+^Ly6G^-^ cells as eosinophils, CD117^+^ cells as mast cells, and CD3^+^ cells as T cells.

### Multiplex cytokine ELISA

Bladder and kidney homogenates were filtered through 0.65 μm Ultrafree®-MC Centrifugal Filter (Millipore sigma). The total protein concentration in organ homogenates was estimated using Pierce BCA protein assay kit (Thermo-scientific). Cytokine levels of IL-1β, IL-6, IL-10, IL-17A, TNF-α, CXCL1 (KC), CCL2 (MCP-1), CCL3 (MIP1α), CCL5 (RANTES), and IFN-γ, in whole bladder and kidney tissue homogenates were estimated using MILLIPLEX® Mouse Cytokine/Chemokine Magnetic Bead Panel (MCYTOMAG-70K-10C). The amount of each cytokine in an individual mouse tissue was presented as a scatter plot showing the amount of cytokine/g of total protein with mean as the measure of central tendency.

### Statistical analysis

The data were analyzed using GraphPad Prism 10.0.0. Data from multiple biological replicates with two or more technical replicates for each experiment were pooled together. The bacterial organ burden and flow cytometry analysis of infiltrating immune cells between different animal cohorts were compared using Mann-Whitney U statistic. The mouse weight, blood glucose levels, and cytokine levels were compared using unpaired t-test. Data were considered statistically significant if P ≤ 0.05.

## RESULTS

### In a mouse model of ascending UTI, DM affects GBS organ burden in a sex-dependent manner

To address our hypothesis that DM exacerbates GBS-UTI, we experimentally induced ascending UTI by transurethral catheterization of GBS 10/84 into the urinary bladders of 8 weeks old, male and female, *db/db* obese, hyperglycemic (diabetic-D), and non-diabetic (ND) mouse littermates. The organ burden was examined in the D and ND littermates from Dock7 background and in D mice alone from BKS background.

At the time of infection, compared to their ND littermates, D-mice were two times heavier (average weight, D = 45.13 ± 3.7 g; ND = 21.6 ± 1.5 g; P < 0.0001; unpaired T test) with two times higher average blood glucose (D= 377 ± 58.8 mg/dL; ND = 166.4 ± 45.6 mg/dL; P < 0.0001; unpaired T test). Within the D cohort, female and male mice showed similar average weights (female= 45.34 ± 5.2 g; male= 44.97 ± 2.3 g; NS) and similar average blood glucose levels (female= 391 ± 67.1 mg/dL; male= 358 ± 40.1 mg/dL; NS). Although, within the ND cohort, compared to females, the male mice showed modestly higher weight (female = 20.5 ± 0.6 g; male = 23.11 ± 1 g; P < 0.0001) and blood glucose levels (female= 149 ± 32.5 mg/dL; male= 219 ± 39.6 mg/dL; P < 0.0001).

Compared to their ND counterparts, the D-mice showed 2000-fold higher median CFU in the urinary bladder (D = 2 × 10^5^ CFU/ml; ND = 100 CFU/ml; P < 0.0001; Mann-Whitney U test; Fig S1A). When stratified for sex, D-male and D-female mice showed higher bladder CFUs than their ND counterparts, although the bladder CFU burden in D-females was substantially higher than that in D-males (Fig 1A): the median bladder burden was ∼10^5^-fold higher in D-females compared to ND-females (D-female = 1.5 × 10^6^ CFU/ml; ND-female = 110 CFU/ml; P < 0.0001) and 14-fold higher in D-males compared to ND-males (D-male = 1000 CFU/ml; ND-male = 70 CFU/ml; P= 0.0651); the median bladder GBS burden in ND-female and ND-male cohorts was similar. In contrast, ∼3-fold increase in the median CFU in D-mice (D = 2 × 10^5^ CFU/ml; ND = 6 × 10^4^ CFU/ml; P = 0.064; Fig S1B) is primarily due to 20-fold higher kidney GBS CFUs in D-males than that in ND-males (D-male = 6 × 10^6^ CFU/ml; ND-male = 3 × 10^5^ CFU/ml; P= 0.0006; Fig 1B); DM did not affect kidney GBS burden in female mice (D-female = 1.5 × 10^4^ CFU/ml; ND-female = 1.7 × 10^4^ CFU/ml; NS ; Fig 1B). The kidney GBS burden in ND-males was 17-fold higher than that in ND-females. These data suggest that DM mediated increase in GBS burden in the UT is sex-dependent.

To study the effects of DM and sex on GBS dissemination resulting in uBSI, we enumerated spleen CFUs in male and female, D and ND mice at 24 hpi. GBS disseminated to the spleens of all (31/31) D-mice and only 2/19 ND-mice (D = 1000 CFU/ml, ND < LOD; P<0.0001; Fig S1C). When stratified for sex, the spleens of D-male and D-female mice exhibited similar median CFUs (D-female = 2000 CFU/ml, D-male = 1000 CFU/ml) and GBS CFUs were above the limit of detection in 2/7 ND males and 0/12 ND female mouse spleens (Fig 1C). These data suggest that DM induces GBS dissemination in a sex-independent manner. Interestingly at 24 hpi, we did not detect CFUs in the blood collected by cardiac puncture of D-mice.

### DM affects immune cell recruitment to the GBS-infected UT in a sex-dependent manner

We performed differential flow cytometry analysis of bladder and kidney tissues from GBS-infected and uninfected (PBS control), D and ND, male and female mice. The scatter plots (Fig 2) represent immune cell counts from bladder (Fig 2 A to E) and kidney (Fig 2 F to J) homogenates from individual mice with median as the measure of central tendency. The flow cytometry data were compared by Mann-Whitney U test.

We noted that (i) the median CD45^+^ total leukocyte in GBS-infected D-bladders was significantly lower (P = 0.0275) than that in GBS-infected ND-bladders (Fig 2A), (ii) the neutrophil counts in the bladder were not affected by either GBS infection or diabetes (Fig 2B); (iii) compared to their uninfected counterparts, GBS infected ND-bladders recruited significantly higher CD45^+^MHCII^-^CD11b^+^SiglecF^+^Ly6G^-^ eosinophils (P = 0.0308; Fig 2C) and CD45^+^CD3^+^ T cells (P = 0.0275; Fig 2E), while (iv) the presence of DM caused a significant reduction in the GBS-mediated recruitment of eosinophils (P < 0.0001; Fig 2C) as well as T cells (P = 0.0007; Fig 2E); and (iv) the GBS-infected D-mice recruit significantly higher CD45^+^CD117^+^ mast cells (P = 0.0245; Fig 2D) to their bladders compared to their ND counterparts.

Furthermore, (i) compared to the uninfected controls, GBS infection induced significant CD45^+^ total leukocyte recruitment in ND-(P < 0.0001) and D-(P = 0.0008) kidneys; (ii) DM significantly reduced the median CD45^+^ leukocyte in both uninfected (P = 0.003) and infected (P < 0.0001) cohorts (Fig 2F). (iii) These changes in total leukocyte recruitment correlated predominantly with the changes in the T cell recruitment as GBS infection significantly induced T cell recruitment to ND-kidneys (P < 0.0001) and the presence of DM reduced GBS-mediated T cell recruitment by ∼6-fold (P < 0.0001) (Fig 2J). (iv) In addition, while GBS-UTI also induced significantly higher median neutrophils in ND-(P = 0.024) and D-(P = 0.0058) kidneys (Fig 2G), higher median eosinophils in D-(P = 0.0096; Fig 2H) kidneys, and higher median mast cells in ND-(P = 0.0171) and D-(P = 0.026) mice (Fig 2I), the presence of DM did not reduce GBS-induced recruitment of these granulocytes to kidneys.

When stratified for sex, GBS-infected ND-males showed modest but significant increase in the median neutrophil compared to GBS-infected ND-female bladder (Fig 2B) and kidneys (Fig 2G).

### DM affects cytokine production in the GBS-infected UT in a sex-dependent manner

Next, we used multiplex ELISA to examine the levels of cytokines (IL-1β, IL-6, IL-10, IL-17, CXCL1, CCL2, CCL3, CCL5, TNFα, and IFNγ) in bladder and kidney homogenates from GBS-infected male and female, D and ND littermates. Compared to their ND counterparts, GBS-infected D-bladders produced significantly lower CXCL1 (P = 0.0165, unpaired t test; Fig.3 A) and significantly higher CCL2 (P = 0.0234; Fig.3 B). The levels of CXCL1 and CCL2 were unaffected by sex while the levels of other cytokines were unaffected by either DM or sex.

Compared to their ND counterparts, GBS-infected D-kidneys produced significantly lower levels of CCL3 (P = 0.0172; Fig.3 C), IL-1β (P = 0.0037; Fig.3 D), and IL-17 (P = 0.0413; Fig.3 E). Notably, ND-males produced significantly higher kidney IL-17 compared to ND-females in response to GBS-UTI (Fig.3.4.B). Overall, the ELISA results indicate that diabetes dampens GBS-induced cytokine production in bladder as well as kidneys.

## DISCUSSION

This study was designed to examine the effects of DM on the pathophysiology of GBS-UTI. DM is a major risk factor for UTI and their exacerbations (27). The inclusion of both male and female mice facilitated, quite serendipitously, sexual dimorphism in GBS urinary tissue tropism. As the model uropathogen, we used GBS 10/84, a hyperhemolytic clinical isolate that belongs to serotype V, one of the three predominant GBS serotypes known to cause UTI in humans (12). We inoculated GBS 10/84, via transurethral catheterization, to induce ascending UTI in the *db*/*db* mouse model of genetically induced DM. Due to a homozygous spontaneous mutation in *Lepr* (leptin receptor), the *db/db* mice exhibit polyphagia which in turn results in obesity, hyperglycemia, and glycosuria around 4-6 weeks of age (28). For our experiments, we bred mice heterozygous for *Lepr* mutation (*db*/*+*) to generate diabetic (D-*db*/*db* homozygotes) and non-diabetic (ND-*db*/*+* or +/+) littermates. The F2-generation littermates are considered a gold-standard for standardizing gut microbiota in experimental animals as they experience the same microbial and environmental exposures up to and following birth (29).

We observed that at 24 hpi, compared to their ND-counterparts, the *db/db* mouse bladders and spleens exhibited significantly higher GBS burden while the *db/db* kidneys showed a substantial yet statistically insignificant increase in GBS CFUs. These observations could be attributed to DM-mediated immune dysfunction marked by the suppression of pro-inflammatory cytokines and CD45^+^ leukocyte recruitment to *db/db* bladder and *db/db* kidney tissues in response to GBS infection. In addition to establishing DM as a risk factor for GBS-UTI, our observations revealed a novel and major role for DM in promoting GBS dissemination from the UT to spleen at 24 hpi. The inclusion of both male and female littermates allowed us to stratify these data and reveal novel sex-specific UT tissue tropism wherein female D-mice were more susceptible to GBS cystitis while the higher susceptibility of males to GBS pyelonephritis was further increased by DM. Both male and female D-mice were equally susceptible to GBS dissemination. Moreover, compared to their female counterparts, both D-as well as ND-male bladders infected with GBS showed significantly higher levels of CXCL1, a neutrophil chemoattractant cytokine, while increased neutrophil recruitment was noted in the bladder as well as kidneys of ND-males with GBS infection.

In 2020, Patras *et al* reported results from a study examining GBS-UTI pathogenesis in the mouse model of STZ-induced DM and their ND counterparts (17). Our observations complement this seminal report such that compared to their ND-counterparts both *db/db* and STZ-mice show significantly higher bladder GBS burden and higher mast cell recruitment to bladder in response to GBS infection. Whether the spleen GBS burden is affected in STZ-mice and whether there are differences in GBS-UTI pathogenesis between male/female, STZ-D/ND mice has not been reported. The differences between the results from STZ and *db*/*db* mice such as the substantial difference in the magnitude of increment in bladder CFUs (1.13-fold for STZ and 2000-fold for *db/db*), different effects of DM on kidney CFUs (DM reduced kidney CFUs in STZ-mice while increasing it in *db/db* mice) may be likely due to the differences in the mouse models of DM (chemically induced versus genetic origin) or GBS strains (COH1 versus 10/84).

Our ongoing work comparing GBS-UTI pathogenesis between D- and ND-littermates at early (6 hpi) and late (days 7 and 28 pi) time points aims to address various acknowledged limitations of this study. For example, (i) blood CFU analysis at 6 hpi will help us determine whether DM promotes transient bacteremia secondary to GBS-UTI early in the infection and may explain why we detected GBS CFUs only in the spleens and not in the blood of all D-mice at 24 hpi, (ii) flow cytometry and ELISA of GBS-infected bladder and kidney tissues from male/female, D/ND mice at 6 hpi will identify immune effectors differentially affected early in the infection by DM and/or sex, (iii) comparing GBS-UTI in male/female, D/ND mice at days 7, 14, and 28 pi will help us the assess the effects of DM and/or sex on chronicity and severity of GBS-UTI complications. Future work is also needed (iv) to examine the pathogenesis of GBS strains other than 10/84 and (v) to explore the differential contributions of obesity, hyperglycemia, and sex to the pathogenesis of UTI caused by different uropathogens including GBS.

## SUPPLEMENTARY FIGURE LEGEND

**Fig S1: The effects of DM on GBS organ burden in the mouse model of GBS-UTI.** Unsegregated CFU data from both male and female littermates within obese, hyperglycemic (closed circles) or their non-diabetic (open circles) cohorts collected at 24 hpi are shown for (A) the urinary bladder, (B) the kidneys, and (C) the spleen. The scatter plots with median as the measure of central tendency and dotted line indicating the limit of detection (LOD) at 10 CFU/ml were compared using Mann-Whitney U test. * indicates P<0.05, **, P<0.01, ***, P<0.001, and ****, P<0.0001.

